# Massively parallel transposon mutagenesis identifies temporally essential genes for biofilm formation in *Escherichia coli*

**DOI:** 10.1101/2020.12.14.409862

**Authors:** Emma R Holden, Muhammad Yasir, A Keith Turner, John Wain, Ian G. Charles, Mark A Webber

## Abstract

Biofilms complete a life cycle where cells aggregate, grow and produce a structured community before dispersing to seed biofilms in new environments. Progression through this life cycle requires temporally controlled gene expression to maximise fitness at each stage. Previous studies have largely focused on the essential genome for the formation of a mature biofilm, but here we present an insight into the genes involved at different stages of biofilm formation. We used TraDIS-*Xpress*; a massively parallel transposon mutagenesis approach using transposon-located promoters to assay the impact of disruption or altered expression of all genes in the genome on biofilm formation. We determined temporal differences in the importance of genes in *E. coli* growing as a biofilm on glass beads after 12, 24 and 48 hours. A selection of genes identified as important were then validated independently by assaying biofilm biomass, aggregation, curli production and adhesion ability of defined mutants. We identified 48 genes that affected biofilm fitness including genes with known roles and those not previously implicated in biofilm formation. Regulation of type 1 fimbriae and motility were important at all time points. Adhesion and motility were important for the early biofilm, whereas matrix production and purine biosynthesis were only important as the biofilm matured. We found strong temporal contributions to biofilm fitness for some genes including some where expression changed between being beneficial or detrimental depending on the stage at which they are expressed, including *dksA* and *dsbA*. Novel genes implicated in biofilm formation included *zapE* and *truA* involved in cell division, *maoP* in DNA housekeeping and *yigZ* and *ykgJ* of unknown function. This work provides new insights into the requirements for successful biofilm formation through the biofilm life cycle and demonstrates the importance of understanding expression and fitness through time.

## Introduction

Bacteria rarely exist planktonically outside of the laboratory and are usually found as part of structured, aggregated communities called biofilms ^1^. Clinically, approximately 80% of infections have been suggested to have a biofilm component ^2^. Biofilm related infections are complicated by their intrinsic tolerance to antimicrobials, making infections difficult to treat and, often persistent ^3–6^. Cells within a biofilm grow more slowly than those in planktonic culture and this reduced level of metabolic activity has been associated with tolerance to antimicrobials, allowing biofilms to be typically 10-1000-fold less sensitive to antibiotics than corresponding strains in planktonic conditions ^7,8^. Aside from the clinical setting, there are many useful applications of biofilms, including wastewater treatment and bioprocessing ^9^. Biofilms undergo a life cycle that commonly consists of initial attachment to a surface, growth and maturation of the biofilm over time with characteristic production of extracellular matrix components, followed by dispersal of planktonic cells to facilitate colonisation of new surfaces ^10^. The switch between planktonic and biofilm lifestyles is driven by environmental stimuli promoting large scale changes in gene expression and regulation that are necessary to support the bacterial community through the life cycle, which is distinct to planktonic growth conditions. Expressing the right genes at the right time and place is critical for efficient production of a biofilm.

The main components of the biofilm extracellular matrix in *E. coli* are the amyloid protein curli, the polysaccharide cellulose and extracellular DNA ^11^. Curli is transcribed by the divergent operons *csgBAC* and *csgDEFG*, with curli biosynthesis regulated by CsgD ^12^. Cellulose biosynthetic machinery is encoded by *bcsRQABZC* and *bcsEFG*, and its production is regulated by c-di-GMP ^13^. Several genes are known to be involved in the regulation of matrix production, including *ompR* ^14,15^, *cpxR* ^14,16,17^ and *rpoS* ^18,19^, amongst others ^20–22^. Extracellular DNA is also an important component of the biofilm matrix, and the addition of DNase has negatively affected the biomass of biofilms formed by *Pseudomonas aeruginosa* ^23^, *Bacillus cereus* ^24^ and a range of Gram-negative pathogens, including *E. coli* ^25^.

Many previous studies have focused on identifying the genes and pathways required for biofilm formation in *E. coli* in the mature biofilm rather than dissecting events across the life cycle. Various studies have taken a genome wide approach to identifying genes that impact biofilm formation. One assessed biofilm formation of all the mutants in the Keio collection ^26^, although analysis is limited to the effect of inactivation ^27^. Another used a transcriptomic approach to identify genes with altered expression in biofilms over time ^28^ and DNA microarrays have also been used to link the presence of different genes with biofilm capacity in panels of isolates ^29^.

Large scale transposon mutagenesis experiments represent another high-throughput, sensitive whole genome approach to link phenotype to genotype methodologies ^30–32^. These methods make use of a massive library of transposon mutants, containing multiple insertions per gene and guaranteeing the presence of a large number of independent deletion mutants of every gene in the genome are represented in the pool. This provides great power in assaying the role of genes as in a high-density library intragenic essentiality of domains with the proteins encoded by genes can also often be inferred by analysing the fitness of multiple independent mutants within a gene. Transposon mutagenesis approaches have however been historically limited by an inability to assay essential genes within which transposon insertions are not viable. This issue has been addressed with our recent development of TraDIS-*Xpress*, which uses transposons containing an outward-facing inducible promoter. Upon addition of an inducer of the transposon-encoded promoter this results in overexpression of genes downstream of transposon insertions, or repression of genes where the transposon is positioned downstream but in an antisense orientation. Therefore, we are able to assay the impact of altering expression of all genes (including those which cannot be inactivated) as well as capturing traditional essentiality measurements, we recently demonstrated the utility of this approach in identifying roles for essential genes in survival of drug exposure ^33^. In this work, we sought to investigate biofilm formation using TraDIS-*Xpress* to get a more detailed view of important genes than possible in the previous studies described above. DNA extracted from biofilms formed by the transposon mutant library on glass beads were directly compared to DNA from planktonic cultures at three time points through the biofilm life cycle. The location and frequency of transposon insertion sites was determined by sequencing and mapped to a reference genome. Differences in insertion frequency between biofilm and planktonic conditions identified a difference in fitness; for example, a gene was considered important for biofilm formation if it contained fewer insertions in biofilm conditions relative to the planktonic control. Predictions made by this approach were then validated using defined mutants from the Keio library, a collection of single knockout mutants in the same parent strain as the transposon mutant library.

This study identified 48 genes that were found to be important at different stages of biofilm formation by *E. coli*. By investigating the genes important across the biofilm life cycle, we were able to get a dynamic view of the main pathways important at different stages of biofilm development. Our findings reinforced the importance of adhesion, motility and matrix production in the biofilm, and revealed the importance of genes not previously implicated in biofilm formation. This included genes with roles in cell division (*zapE* ^34^ and *truA* ^35^), DNA housekeeping (*maoP* ^36^) and *yigZ* and *ykgJ*, the functions of which have not been elucidated. We identified clear requirements for some pathways at specific points of the biofilm life cycle, furthering our understanding of how biofilms maintain fitness over time.

## Results

### Confirmation of model efficacy

Wild type *E. coli* BW25113 was grown on glass beads and harvested over time to investigate biofilm development after 12, 24 and 48 hours (figure 1). The changes in biofilm CFU (figure 1a) and architecture (figure 1b) after 12-, 24- and 48-hours growth demonstrate different phases of biofilm formation. A transposon mutant library containing approximately 800,000 unique mutants was then grown on glass beads and harvested at the same time intervals. The genomic DNA obtained from biofilms and planktonic culture each time point was analysed following the TraDIS-*Xpress* methodology to determine the differences in gene essentiality and expression over time. This found 48 genes that considerably affected biofilm formation over time in *E. coli*: 42 were identified as being needed to maintain the fitness of a biofilm and expression of 6 genes was predicted to result in reduced biofilm fitness (figure 2 and supplementary table 1). The main pathways that were consistently important in the biofilm through all the time points included type 1 fimbriae, curli biosynthesis and regulation of flagella. All other loci identified affected biofilm formation at specific points in the life cycle.

**Figure 1:**
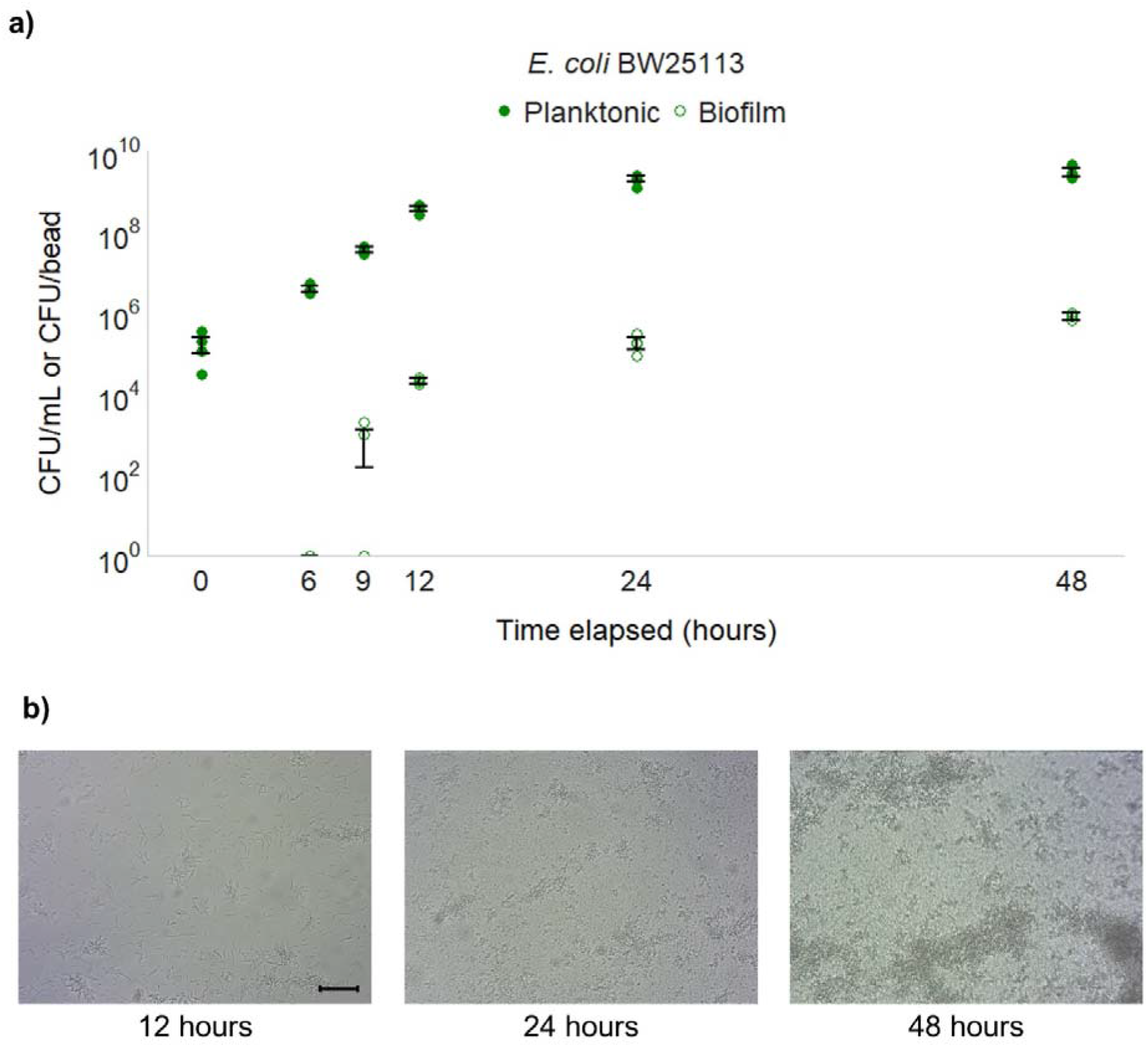
Biofilm formation of wild type *E. coli* BW25113 on glass over time. a) CFU of planktonic and biofilm samples harvested from the model at different time points through biofilm development. Planktonic samples are measured in CFU per mL of culture and biofilm samples are measured in CFU of cells isolated from one glass bead. Points represent four independent replicates and error bars show 95% confidence intervals where present. b) Biofilms formed on glass under flow conditions after 12, 24 and 48 hours growth. Images are representative of two independent replicates. 20x Magnification. Scale bar indicates 5 μm.

**Figure 2:**
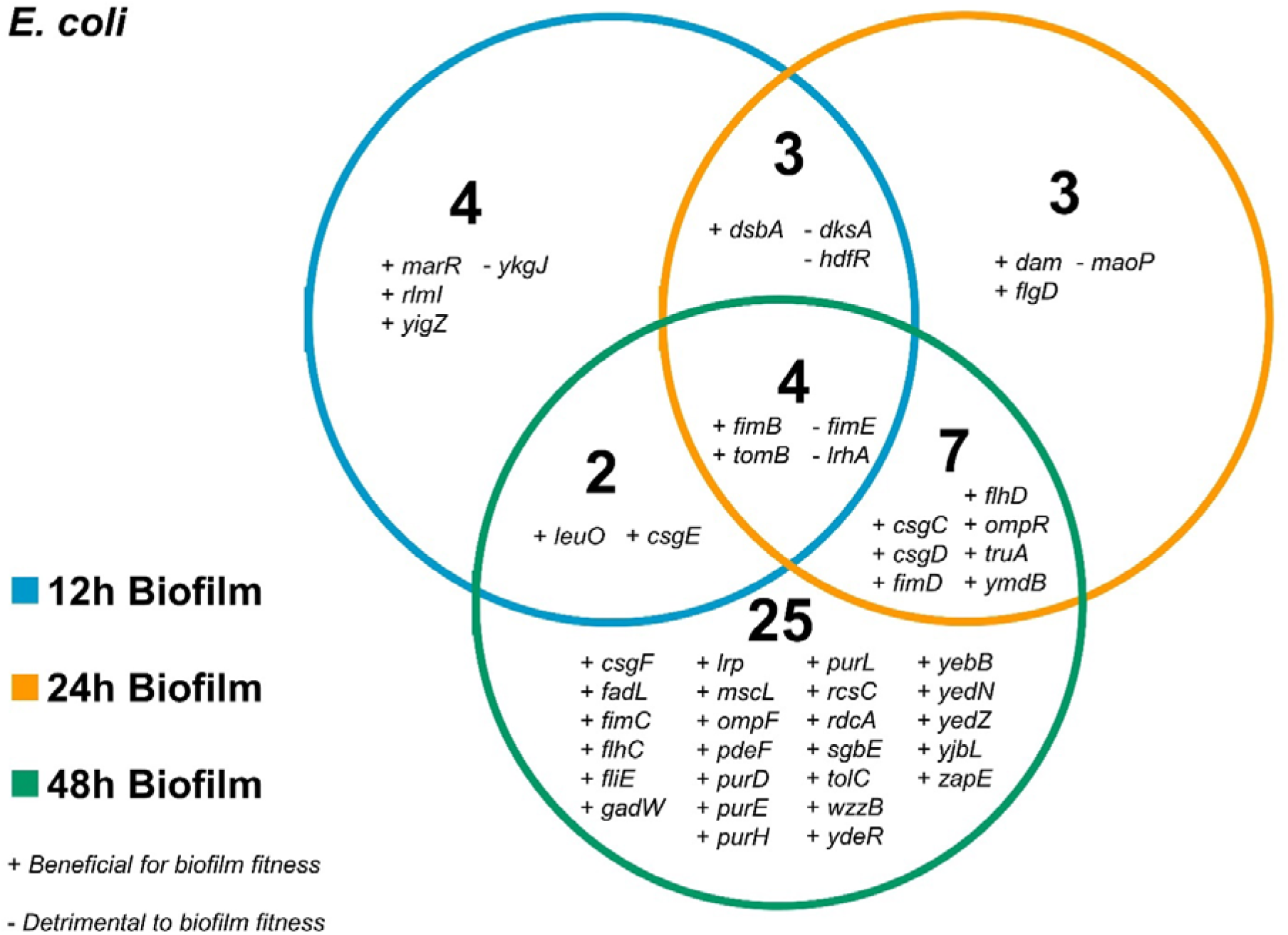
Genes involved in biofilm formation over time in *E. coli*. Plus (+) symbols indicate genes that were beneficial for, and minus (−) symbols indicate genes that were detrimental to, biofilm formation.

### Fimbriae expression and motility are important at all stages of biofilm formation

Only 4 genes were found to be important throughout 12, 24 and 48 hours (figure 2). These included *fimB* and *fimE*. The recombinase gene *fimB* which helps mediate both ‘ON-to-OFF’ and ‘OFF-to-ON’ switching of fimbriae expression was beneficial for biofilm formation at all time points. Fewer *fimB* mutants were observed in biofilm conditions compared to planktonic, and this number decreased over time. In contrast, inactivation of *fimE*, responsible for only ‘ON-to-OFF’ fimbrial regulation ^37^, increased biofilm fitness at all time points. Initially, there were only slightly more *fimE* mutants in biofilm conditions compared to planktonic at 12 hours, but this increased over time with a stark contrast seen between biofilm and planktonic conditions at the 24- and 48-hour time points (figure 3a). The predicted impacts on biofilm formation were phenotypically validated by testing both gene knockout mutants from the Keio collection (which contains two, independent deletion mutants for most genes in *E. coli* BW25113) ^38^. Biofilm biomass was measured by growing knockout mutants in a 96-well plate for 48 hours and staining the resulting biofilm with crystal violet. Cell aggregation was quantified by measuring the optical density of the supernatant of cultures left unagitated for 24 hours. Analysis of Δ*fimB* and Δ*fimE* mutants showed no deficit in biofilm biomass (figure 4a), but both were deficient in cell aggregation, determined by an increased optical density of stagnant culture supernatants relative to wild type cultures (figure 4b). Together, the TraDIS-*Xpress* and phenotypic data suggest that the ability to regulate fimbriae expression in a phase-dependent manner is important for fitness of a biofilm, rather than being constrained in a fimbriae ‘ON’ or ‘OFF’ state.

**Figure 3:**
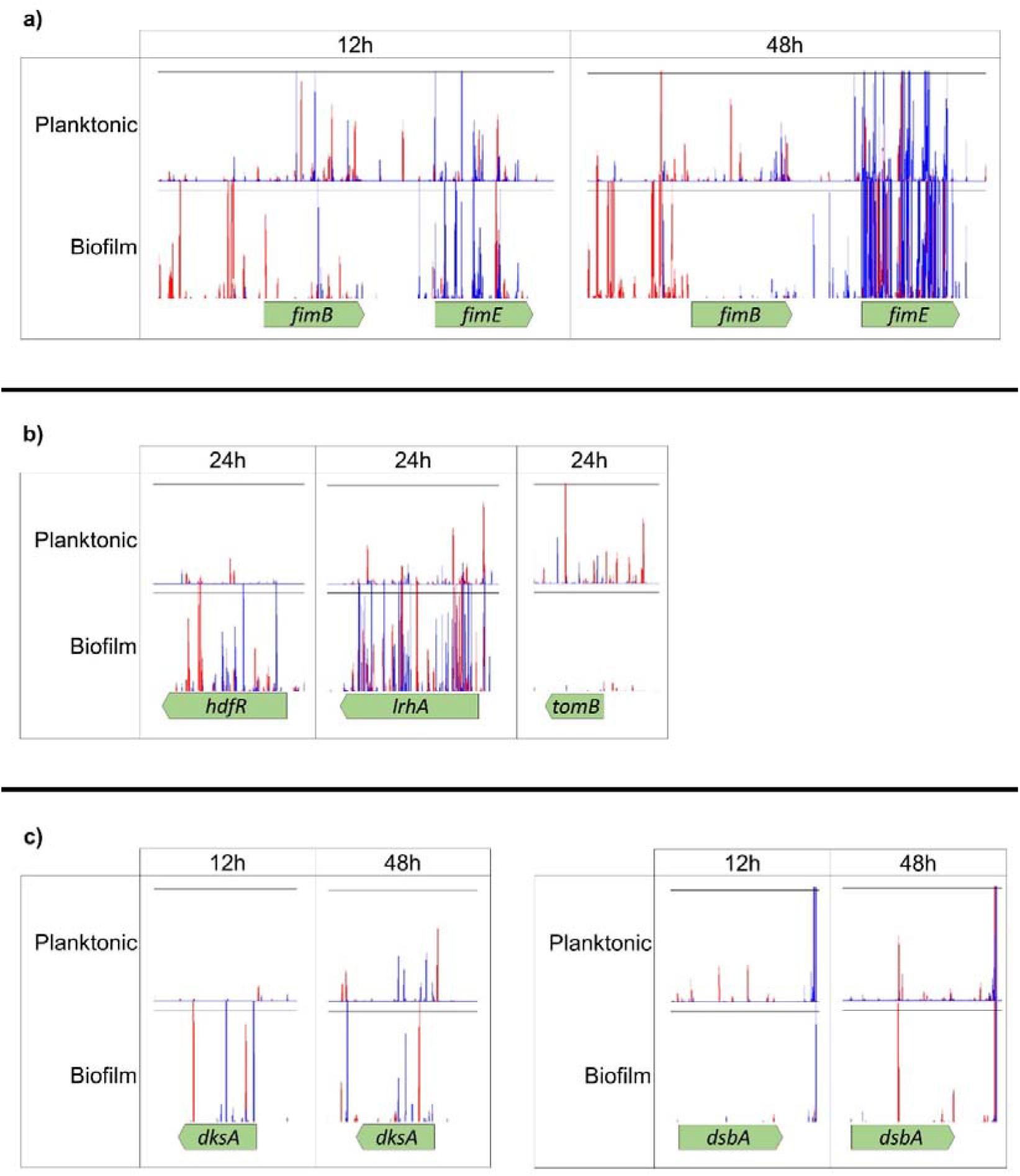
Transposon insertion sites and frequencies in planktonic and biofilm conditions, mapped to a reference genome and plotted with BioTraDIS in Artemis. The height of the peak can be used as a proxy for the mutant’s ‘fitness’ in the condition. Red peaks indicate where the transposon-located promoter is facing left-to-right, and blue peaks show it facing right-to-left. **a)** Insertion sites in and around *fimB* and *fimE* in planktonic and biofilm conditions after 12- and 48-hours growth. **b)** Insertion sites in and around *hdfR*, *lrhA* and *tomB* in planktonic and biofilm conditions after 24 hours growth. **c)** Insertion sites in and around *dksA* and *dsbA* in planktonic and biofilm conditions after 12- and 48-hours growth. For all plot files, one of two independent replicates is shown, and the transposon-located promoter is induced with 1mM IPTG in all conditions shown.

**Figure 4:**
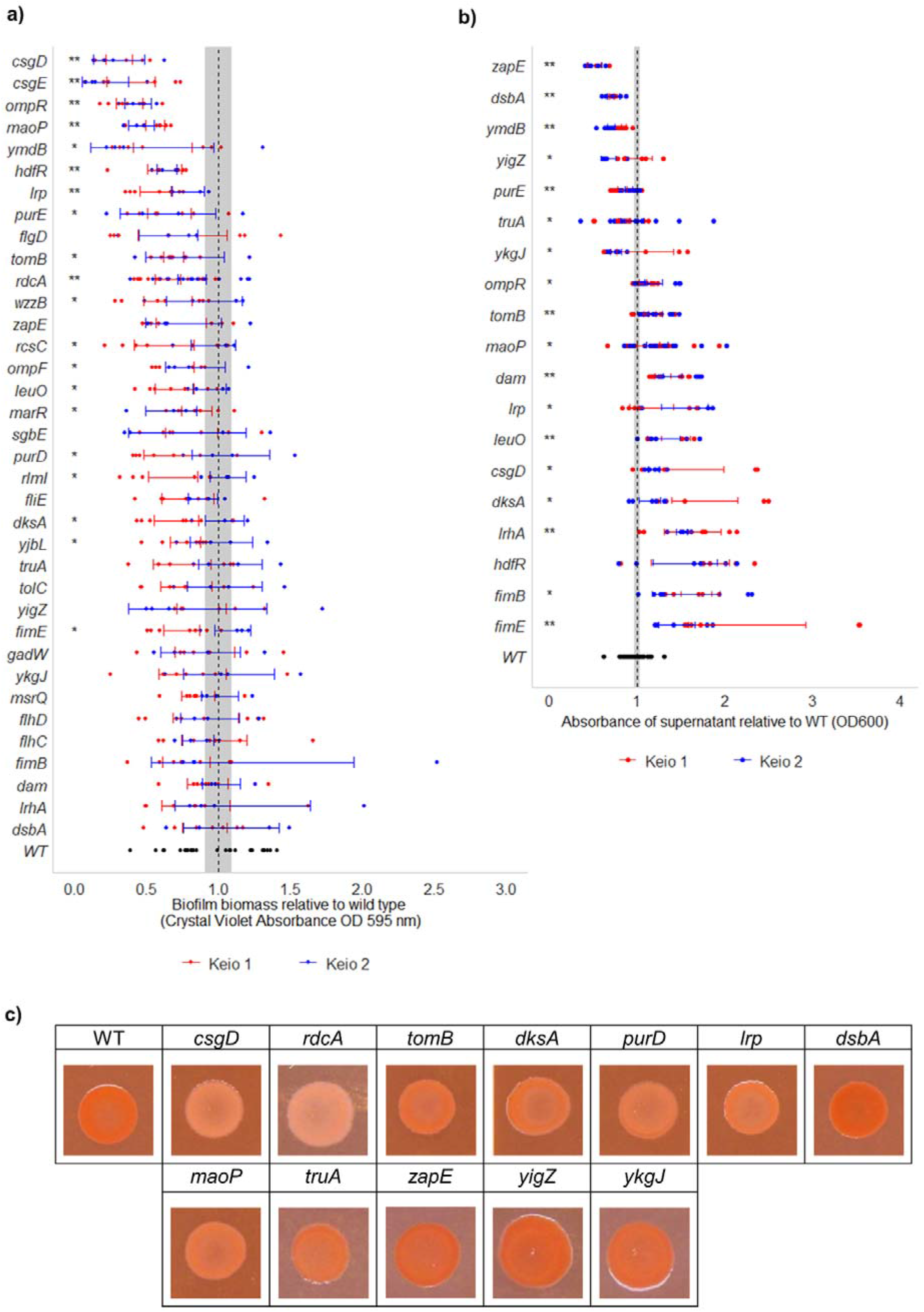
Phenotypic validation of selected genes involved in biofilm formation. **a**) Biofilm biomass of single knockout mutants relative to wild type *E. coli*, measured by crystal violet staining. Two biological and a minimum of two technical replicates were performed for each mutant. **b**) Cell aggregation of single knockout mutants relative to wild type *E. coli*, measured by OD_600 nm_ of the supernatant of unagitated cultures. Points show the ODs of three independent replicates. For both graphs, red points/bars distinguish between the two Keio collection mutants of each gene. Error bars show 95% confidence intervals, and the shaded area shows the 95% confidence interval of the wild type. Single asterisks (*) represent a significant difference between one Keio mutant copy and the wild type, and double asterisks (**) denote a significant difference between both Keio mutant copies and the wild type (Welch’s *t*-test, *p* < 0.05). **c**) Colonies grown on agar supplemented with Congo red to compare curli biosynthesis between single knockout mutants and the wild type. Images are representative of 2 biological and 2 technical replicates.

Disruption of *lrhA*, a regulator of motility and chemotaxis ^39^, was beneficial for biofilm formation at all time points (figure 3b). LrhA also has a role in type 1 fimbriae expression through activating expression of *fimE* ^40^, but in addition represses flagella-mediated motility. Analysis of the Δ*lrhA* biofilm showed initial formation of microcolonies occurred faster than the wild-type (figure 5a) but at later time points the biofilms formed by this mutant were less mature than seen with the wild-type. There was no significant change in biomass formed by this mutant (figure 4a) and mutants appeared less aggregative than the wild type (figure 4b). These data suggest that inactivation of *lrhA* impacts both adhesion and aggregation differently at distinct stages of the biofilm life cycle and may result in a benefit to early surface colonisation but with a cost to later maturation.

**Figure 5:**
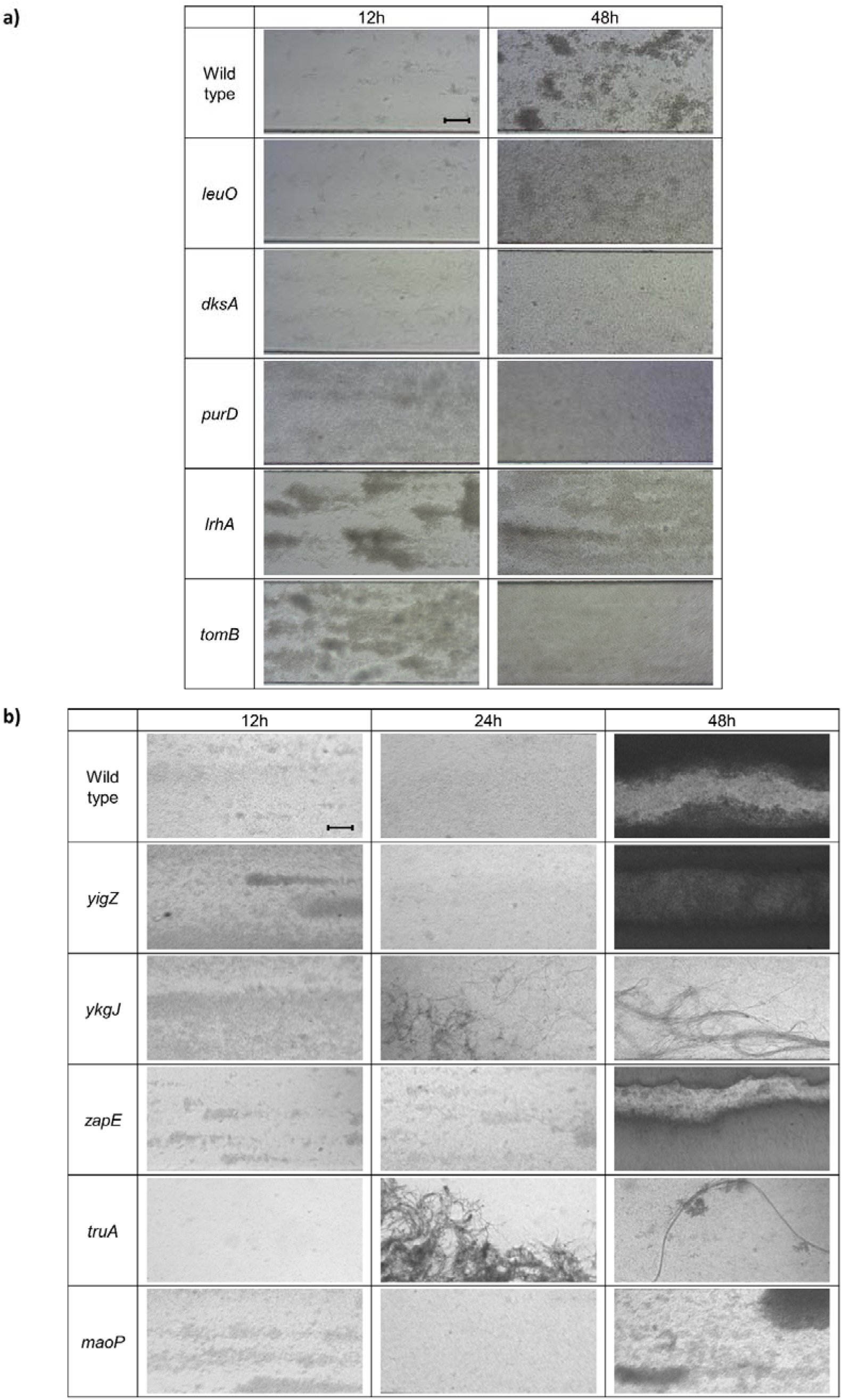
Biofilm formation of single knockout mutants on glass analysed under flow conditions after 12-, 24- and 48-hours growth. a) Single knockout mutants selected for their effect on biofilm fitness. B) Single knockout mutants of genes not previously described to affect biofilm formation, to the best of our knowledge. 10x Magnification. Images are representative of two independent replicates. Scale bar indicates 10 μm.

Expression of the Hha toxin attenuator *tomB* was also found to be consistently important for biofilm formation at 12, 24 and 48 hours (figure 3b). Consistent with this prediction, the Δ*tomB* mutant biofilm had reduced cell aggregation and curli biosynthesis, and reduced biofilm biomass (figure 4 a,b,c). Under flow conditions, the Δ*tomB* mutant biofilm has a similar appearance to the Δ*lrhA* mutant biofilm, with microcolonies visible after 12 hours growth which disappeared over time (figure 5a).

### Regulatory genes are important in the early biofilm

In the early biofilm, at 12 hours growth, only 13 genes were found to distinguish the planktonic and biofilm conditions. TraDIS-*Xpress* data indicated that inactivation of transcriptional factor *dksA* promoted biofilm formation at the 12- and 24-hour time points but not in the mature biofilm (figure 3c). Supporting this, analysis of Δ*dksA* mutant biofilms under flow conditions showed an initial benefit with increased adhesion at both the 12- and 24-hour time points, but reduced microcolony formation at the 48-hour time point, suggesting *dksA* affects biofilm initiation (figure 5a). Inactivation of Δ*dksA* was also seen to reduce cell aggregation, curli biosynthesis and biofilm biomass (figure 4 a,b,c). Expression of *hdfR*, a negative regulator of motility ^41^, was found to be detrimental to biofilm fitness in the early biofilm after 12- and 24-hours growth (figure 3b), and Δ*hdfR* mutant biofilms had significantly reduced biomass (figure 4a). In addition, the stress response regulator *marR* ^42^ and the 23S rRNA methyltransferase *rlmI* ^43^ were both found to be beneficial for biofilm fitness at the 12 hour time point only, and reduced biofilm biomass was found in the corresponding deletion mutants (figure 4a). These genes have both previously been implicated in biofilm formation ^43–45^, but the effect on early biofilm formation has not been described previously.

Two genes of unknown function, *yigZ* and *ykgJ* were found to affect biofilm formation at 12 hours. Fewer mutants were observed in *yigZ* in biofilm conditions relative to planktonic at 12 hours, indicating its importance in early biofilm formation. We also saw that reduced expression of *ykgJ* was beneficial for biofilm formation, with more transposon insertions antisense to *ykgJ* present in biofilm conditions relative to planktonic. Although there were no differences seen between the wild type and *ykgJ* in biofilms grown under flow conditions for 12 hours, differences became apparent at the 24- and 48-hour time points, where the *ykgJ* mutant is significantly more filamented. For both *yigZ* and *ykgJ*, one mutant copy showed slightly increased aggregation relative to the wild type (figure 4b), but there were no differences observed in biofilm biomass, curli biosynthesis or adhesion (figures 4 a, c and figure 5b).

### Biofilms sampled after 24 hours demonstrate both adhesion and matrix production are important

More pathways were identified as being important to biofilm formation at 24 hours that at 12 hours. Two genes involved in DNA housekeeping were found to be involved in biofilm formation at this time point. These included *dam*, encoding DNA methyltransferase ^46^, insertional activation of which was not tolerated in the 24 hour biofilm, with Δ*dam* mutants defective in aggregation compared to the wild type (figure 4b). Also, inactivation of *maoP,* involved in Ori macrodomain organisation ^36^, was predicted to confer a fitness advantage in the 24-hour biofilm compared to the planktonic condition. TraDIS-*Xpress* data showed more reads mapped to *maoP* in the biofilm conditions compared to the planktonic at 24 hours suggesting loss of this gene was beneficial. Phenotypic analysis of the Δ*maoP* mutant biofilm did demonstrate a phenotype although in opposition to the prediction, *maoP* mutants were significantly deficient in biofilm biomass production, curli biosynthesis and one mutant displayed reduced aggregation (figure 4 a, c). After 48 hours growth under flow conditions, Δ*maoP* mutant biofilm was considerably less dense than the wild type (figure 5b).

In the 12- and 24-hour biofilms, *dsbA* (encoding disulphide oxidoreductase ^47^) was essential with no insertions detected within this gene (figure 3c). The role of *dsbA* in adhesion to abiotic surfaces and epithelial cells has previously been suggested ^47,48^. Phenotypic validation of the Δ*dsbA* mutant showed a red, dry and rough (*rdar*) phenotype on Congo red plates (figure 4c), indicative of increased curli biosynthesis. Cell aggregation in the Δ*dsbA* mutant was significantly improved compared to the wild type, implying a role of *dsbA* in inhibiting cell-cell aggregation. Our data showed that *dsbA* is important in the early biofilm, but its deletion appears to be beneficial to the formation of a mature biofilm, according to the Congo red and aggregation data. Supporting this hypothesis, *dsbA* was not essential at the 48-hour time point.

Inactivation of the RNase III modulator *ymdB* ^49^ was found to reduce fitness in the biofilm, with fewer reads mapping here in biofilm conditions compared to planktonic at both the 24- and 48-hour time points. This follows previous findings that both inactivation and overexpression of *ymdB* negatively affects biofilm biomass ^49^. In concordance with these findings, a Δ*ymdB* mutant had significantly improved cell aggregation and reduced biofilm biomass (figure 4 a,b).

Curli biosynthesis became important by the 24 hour time point as no insertions mapped to *csgC*, encoding a curli subunit chaperone ^50^ and more transposon insertions mapped upstream of the curli biosynthesis regulator *csgD* ^12^, suggesting its increased expression benefitted biofilm formation. At the 48 hour time point, both genes were essential for biofilm formation, which was also the case for the known *csgD* regulator, *ompR* ^14^. This importance was supported by significantly reduced biofilm biomass and reduced aggregation in knockout mutants (figure 4a, b).

### The mature biofilm grown for 48 hours requires purine biosynthesis, matrix production, motility and solute transport

There were 38 genes found to be important for fitness of the mature biofilm after 48 hours growth, and 25 of these genes were identified as essential at this time point only. The major pathway implicated in biofilm formation at 48 hours was purine ribonucleotide biosynthesis, with four genes, *purD*, *purH*, *purL* and *purE* ^51^, found to be essential at this time point only. TraDIS-*Xpress* did not identify mutants in any of these genes in biofilms sampled at 48 hours, whereas several reads mapped to these loci under planktonic conditions, as well as under both biofilm and planktonic conditions earlier at 12 and 24 hours. Visualisation of a Δ*purD* mutant biofilm under flow conditions saw poor biofilm formation and no microcolony formation at any time compared to the wild type (figure 5a). Additionally, Δ*purD* and Δ*purE* mutants were deficient in biofilm biomass production, curli biosynthesis, and Δ*purE* also showed increased cell aggregation (figure 4 a,b,c), confirming an important role for purine biosynthesis in matrix production and curli biosynthesis in the mature biofilm.

The flagella master regulatory system *flhDC* was identified as important in the mature biofilm. Biofilms sampled after 48 hours saw fewer *flhC* mutants, while insertions interpreted as over-expressing *flhD* increased in numbers both at the 24- and 48-hour time points, compared to planktonic conditions. No mutants in *flgD* and *fliE*, encoding flagellar filament proteins, were identified at 24 and 48 hours, respectively. It has previously been shown that motility is important for initial biofilm formation ^52,53^, but this may not relate to biomass formation where no differences were seen for Δ*flhD*, Δ*flhC*, Δ*fliE* and Δ*flgD* mutants.

Various pleiotropic transcriptional regulators were also important in the mature biofilm. This included the H-NS antagonist *leuO* ^54^. Increased insertions upstream of *leuO* under biofilm conditions after 12 hours growth, as well as no *leuO* mutants in 48-hour biofilms, indicated it was beneficial to biofilm formation. A Δ*leuO* mutant did not aggregate as well as the wild type, and one Δ*leuO* mutant had reduced biofilm biomass (figure 4 a,b). The Δ*leuO* mutant biofilm under flow conditions demonstrated an inability to form microcolonies after 48 hours growth (figure 5a). The leucine-responsive global regulator *lrp* ^55^ and a transcriptional regulator responsible for survival under acid stress, *gadW* ^56^ were also found to have fewer mutants in the 48 hour biofilm compared to the planktonic condition, indicating their importance in the mature biofilm. Reduced biofilm biomass, aggregation and curli biosynthesis were observed for one copy of Δ*lrp*, but no differences in biofilm formation or aggregation were seen for Δ*gadW* mutant biofilms (figure 4 a,b,c).

Inactivation of the outer membrane channels *mscL* ^57^, *tolC* ^58^ and *ompF* ^59^ was not tolerated in the mature biofilm grown for 48 hours. This would indicate the importance of transport in the mature biofilm, however inactivation of *tolC* and *ompF* did not result in a change in biofilm biomass (figure 4a).

Two genes involved in cell division, *zapE* ^34^ and *truA* ^35^, were identified as important in the 48-hour biofilm. No mutants were seen within *zapE* in biofilms grown for 48 hours, indicating its essentiality for biofilm formation at this stage. This was however not reflected in the phenotype of the defined deletion mutants tested, with no changes observed in biofilm biomass or curli biosynthesis, and increased aggregation seen in Δ*zapE* mutants relative to the wild type (figure 4 a, b, c). However, a *zapE* mutant had considerably reduced adhesion after 12 hours growth under flow conditions, relative to the wild type (figure 5b). The pseudouridine synthase *truA* ^60^ was found to be essential in the mature biofilm grown for 24 and 48 hours. When grown under flow conditions, the Δ*truA* mutant cells were extremely filamented in the biofilm compared to the planktonic cells in the same space (figure 5b). After 48 hours, and only a line of filamented cells remains.

## Discussion

We have characterised the essential genome of *E. coli* biofilms across the biofilm lifecycle by using the high throughput transposon mutagenesis screen TraDIS-*Xpress* (figure 6). The identification of genes and pathways already described to be involved in biofilm formation validates the efficacy of this experimental model and shows how assessing many mutants in parallel can identify a large number genes involved in a phenotype. The early biofilm established 12 hours after inoculation was characterised by genes involved in adhesion. The 24-hour biofilm required both adhesion and matrix production, transitioning into matrix production being of the upmost importance in the mature biofilm after 48 hours. In concordance with previous work identifying genes whose importance varies with time in the *E. coli* biofilm, we also reported that control of fimbriae expression and motility remained important at each stage of the biofilm life cycle rather than just being involved in initial attachment ^28^.

**Figure 6:**
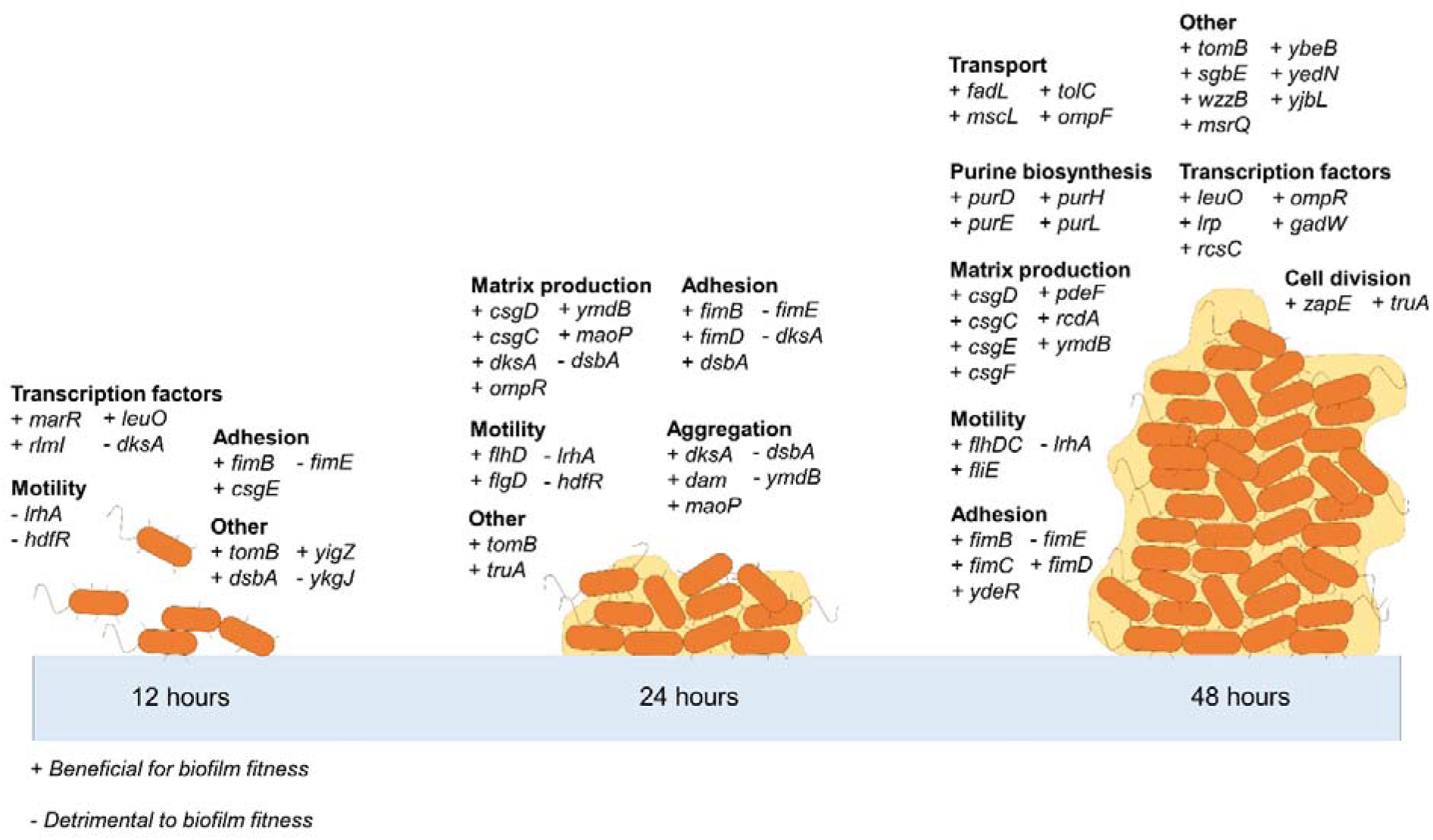
Summary of genes important for biofilm formation by *E. coli* at different stages of development.

As well as identifying how the presence or absence of genes affected biofilm formation, TraDIS-*Xpress* can also determine how increased or reduced gene expression affects biofilm fitness. We found the overexpression of 3 genes and reduced expression of 1 gene was beneficial for biofilm fitness. This could not be done with traditional transposon mutagenesis and provides a further depth to our understanding of how gene expression affects biofilm formation. TraDIS-*Xpress* was also able to identify several genes not previously reported to be involved in biofilm formation, to our knowledge, including *yigZ*, *ykgJ*, *zapE*, *maoP* and *truA*. We found that expression of *maoP* was detrimental to the fitness of biofilms grown for 24 hours, and a Δ*maoP* mutant biofilm had reduced biofilm biomass and reduced curli biosynthesis compared to the wild type. A homolog to *maoP* in *Yersinia pestis* was identified as having a role in adhesion and may positively regulate adhesin expression ^61^. Chromosomal organisation of the Ori macrodomain requires both *maoP* and *maoS* ^36^, however no signal is seen in our data for *maoS*. Further investigation into how chromosomal macrodomain organisation affects biofilm formation is warranted.

The importance of cell division in the mature biofilm was further supported by our findings of fewer *zapE* and *truA* mutants surviving in biofilm conditions compared to planktonic conditions. We found reduced adhesion in the Δ*zapE* mutant biofilm and increased filamentation in the Δ*truA* mutant biofilm. ZapE has been found to be required for growth under low oxygen conditions as well as its role in cell division ^34^, and this may be relevant for why its expression was beneficial for cells within a submerged biofilm. Deletion of *truA* has previously been reported to result in filament formation and reduced cell division ^35^, intracellular survival and oxidative stress conditions ^62^. Deletion of *ykgJ* was also found to cause filament formation in biofilms grown for 24 and 48 hours, which again suggests a role in cell division for this gene. Filamentation has previously been suggested to provide a competitive advantage in adhesion and early biofilm formation, but filamented cells were outcompeted as the biofilm matured ^63^. Our data suggests that temporal regulation of filamentation is important for optimal biofilm fitness over time.

Expression of *dsbA* and repression of *dksA* was found in this study to benefit early biofilm fitness. Based on previous studies and phenotypic analysis of knockout mutants in this study, we believe the increase in biofilm fitness seen is due to increased adhesion in these mutants ^48,64^. This study has highlighted the benefit of close temporal gene regulation in the biofilm, where the expression of certain genes can have a different effect on biofilm fitness at different stages of the biofilm life cycle. We found that *dsbA* was important for the early biofilm where adhesion is crucial and DsbA-DsbB is known to chaperone various adhesins. A *dsbA* mutant biofilm was poor at adhesion although did demonstrate increased curli expression and increased aggregation. Expression of *dsbA* has been previously found to result in repression of the curli regulator *csgD* and curli subunit *csgA*, essential for optimal fitness of the mature biofilm ^65^. Conversely, we found that expression of the transcription factor *dksA* was detrimental in the early biofilm, whilst a *dksA* knockout biofilm had reduced biofilm biomass, reduced curli biosynthesis and reduced aggregation. The effect of *dksA* expression of biofilm formation has been extensively studied and it is known that the deletion of *dksA* increases fimbriae-dependent adhesion, but reduces motility ^64^ and curli production ^32,66,67^. Again, these data show differential expression of important genes at different stages of the biofilm life cycle is essential for optimising biofilm fitness.

Purine biosynthesis was found to be important in the mature biofilm, through the essentiality of *purD*, *purE*, *purL* and *purH* in biofilms grown for 48 hours. Similar findings have previously been described in another transposon mutagenesis experiment in uropathogenic *E. coli* ^32^. Inactivation of purine biosynthetic genes was also found to impair biofilm formation in *Bacillus cereus*, but this was thought to be due to reduced extracellular DNA in the biofilm matrix ^24^. Extracellular DNA is thought to aid adhesion and has been found to be important in the biofilms of a wide range of bacterial species ^23,25^. Our data suggest the importance is in the mature biofilm rather than initial adhesion. A relationship between both purine and pyrimidine biosynthesis and curli production in the biofilm has been reported ^32,66,68^. More recently, curli biosynthesis in a *purL* mutant was reported to be abrogated through addition of inosine, which is involved in the *de novo* purine biosynthetic pathway for production of adenosine monophosphate (AMP) and guanine monophosphate (GMP) ^69^. This suggests that nucleotide production itself, rather than the regulatory effects of the genes involved, affects curli biosynthesis, supporting one hypothesis that disruption of the purine biosynthetic pathway may directly result in a reduction of cyclic-di-GMP. Additionally, we identified two genes involved in c-di-GMP biosynthesis, *rcdA* and *pdeF* ^70^, to be important for biofilm formation at 48 hours. It has been well described how c-di-GMP affects biofilm biomass production and curli biosynthesis ^32,70^. Quantification of intracellular c-di-GMP or further investigation of other c-di-GMP-dependent pathways in these mutants would uncover the relationship between these pathways and biofilm formation.

TraDIS-*Xpress* data suggested that expression of *fimB* and deletion of *fimE* was necessary for optimal biofilm fitness at all points in the biofilm, rather than just for initial attachment. Analysis of Δ*fimB* and Δ*fimE* deletion mutants found no significant change in biofilm biomass and reduced cell-cell aggregation. This finding supports previous work that reported increased fimbriae expression across all stages of biofilm development ^28^. Previous work has observed a positive correlation between type 1 fimbriae and exopolysaccharide production in the mature biofilm ^71^, but the increase in biofilm biomass to support this was not seen in our study. These data suggest that for a population the ability to present cells both with and without fimbriae is beneficial for fitness throughout the life cycle.

Analysis of biofilms under flow conditions found that Δ*lrhA* and Δ*tomB* mutant biofilms had a similar appearance after 12 hours growth, which could indicate a similar role in the biofilm. The role of *lrhA* in motility regulation has been well documented ^39,40,72^, and expression of *tomB* has been seen to reduce motility through repression of *fliA* ^73^. We found deletion of either *lrhA* or *tomB* resulted in reduced aggregation. Although Δ*lrhA* and Δ*tomB* deletion mutants shared many similar phenotypes, TraDIS-*Xpress* data predicted that *tomB* was beneficial and *lrhA* was detrimental to biofilm formation at 12, 24 and 48 hours. Previous studies on Δ*lrhA* mutant biofilms have reported increased adhesion, aggregation and biomass compared to the wild type ^40^. Although we reported decreased aggregation in the Δ*lrhA* mutant after 24 hours (figure 4), we also saw increased adhesion, aggregation and flow cell coverage at each time point in Δ*lrhA* mutant biofilms grown under flow conditions relative to the wild type (figure 5a). This supports the findings from the TraDIS-*Xpress* data, showing inactivation of *lrhA* was beneficial for biofilm fitness throughout biofilm development. This may be due to reduced induction of *fimE* by LrhA ^40^, thereby allowing expression of type 1 fimbriae to facilitate adhesion. We have already described how expression of both *fimB* and *fimE* is necessary for optimal fitness of the mature biofilm, and the effect of *lrhA* on biofilm formation correlates with these findings, with reduced aggregation in Δ*lrhA* biofilms after 24 hours (also seen in *fimB* and *fimE* mutants) and no microcolony formation under flow conditions at 24 and 48 hours. Therefore, the importance of *lrhA* to biofilm formation clearly appears to be time dependent, with the most important role in early events.

Studies on the effect of *tomB* on biofilm formation have focused on its toxin-antitoxin relationship with *hha*, which has been found to reduce expression of fimbrial subunit *fimA* and activate prophage lytic genes causing cell death ^74^. Deletion of *hha* was found to reduce motility through *flhDC* and increase curli production through *csgD* ^75^. We found no obvious benefit to biofilm fitness with insertional inactivation of *hha*, but this may not be visible in our data due to these mutants having a functional copy of *tomB*. We predict *tomB* is involved in regulation of adhesion in the early biofilm and matrix biosynthesis in the late biofilm.

The relationship between motility and biofilm formation is complex. Although it is widely understood that motility is crucial for biofilm formation ^52,53^, it is also true that motility and curli production have an inverse relationship, where biofilm matrix production is induced and motility is repressed for a motile-to-sessile lifestyle transition ^67,77,80^. We found that although insertional inactivation of negative motility regulators *lrhA* and *hdfR* improved biofilm fitness according to the TraDIS-*Xpress* data, a Δ*hdfR* deletion mutant had reduced biofilm biomass, and deletion of either *lrhA* or *hdfR* impaired cell aggregation. Our data found an important role for structural flagella components in the mature biofilm. Previous work has suggested that flagella filaments are important for initial attachment and adhesion ^81^, however we did not find this to be the case, with the expression of flagella filaments only appearing to increase biofilm fitness in the mature biofilm grown for 48 hours. This is supported by previous work that found expression of flagella is important at all stages of the developing biofilm ^28^. It appears that regulation of flagella and motility, rather than their fixed expression or absence, is important for optimal biofilm fitness.

Previous genome-wide screens on *E. coli* biofilm formation have identified many of the same genes as this study ^26,30,32^. The TraDIS-*Xpress* technology used here makes for a more a powerful analysis of biofilm fitness by predicting the effect of changes in gene expression and gene essentiality over time. Differences between this work and previous studies may reflect biofilms being grown under different conditions on different surfaces, as these environmental factors greatly affect the pattern of gene expression and gene essentiality in the biofilm ^82^. Additionally, whole gene knockout mutants, such as those constructed for other studies and those in the Keio collection, differ from transposon insertion mutants. With an insertion an average every 6 base pairs, TraDIS-*Xpress* gives a more in-depth analysis of exactly which regions of the genes in question are important for a given phenotype ^33^. In addition, the TraDIS-*Xpress* experiments involved competition of each mutant against the rest of the pool, this is very sensitive to changes in fitness. Whilst we chose a set of important biofilm-associated phenotypes for validation of our defined mutants these are inevitably somewhat crude and cannot replicate the competition happening within the biofilms in the main experiments. It is likely we failed to identify the basis for a phenotypic impact of some of our candidate mutants in our limited validation conditions with whole gene inactivation mutants.

Various genes were expected to be identified by the model to confirm its efficacy, such as genes involved in curli biosynthesis, however there were some genes that were not detected by TraDIS-*Xpress* that are known to affect biofilm formation. Although many genes involved in curli biosynthesis were identified by our model, the gene encoding the main curli subunit, *csgA*, was not detected. We believe this is because TraDIS-*Xpress* experiments use a mutant library pool, where CsgA produced by the surrounding population can complement the Δ*csgA* mutants ^83^. Although this may be a potential limitation for studying a gene’s role in biofilm formation, is it more representative of intercellular interactions in a non-clonal multispecies biofilm found outside the laboratory. We also did not identify antigen 43 (*agn43/flu*) as important for biofilm formation, despite its strong role in aggregation and adhesion ^84,85^. Previous work found Antigen 43 was important for biofilm formation in glucose-minimal media, but not LB ^85^. This fuels the need for more genome-wide studies analysing a wide range of environmental conditions, strains and species, abiotic and biotic surfaces, to provide a wider list of essential genes for biofilm formation shared amongst important human pathogens, as well as substrate-, condition- and species-specific genes and pathways for specific industrial, clinical and drug-development applications. As well as temporal changes in gene expression, spatial changes have been shown to affect biofilm development ^86^. Integration of the spatial component into this model, to assay how gene expression throughout the biofilm over time affects biofilm fitness, would be the next logical step in furthering our understanding of biofilm development.

This study had revealed important time-specific roles for known and novel genes in biofilm formation. We reveal some pathways have a more important role in the mature biofilm than previously appreciated and identify genes with time dependent conditional essentiality within the biofilm. We also identify potential new candidate genes essential for biofilm formation, which could be targeted for novel anti-biofilm therapies. Further work using high-density transposon mutant libraries across time and in different conditions is likely to further our understanding of biofilm biology.

## Methods

### Transposon mutant library

The *E. coli* BW25113 transposon mutant library containing over 800,000 distinct mutants that was used in this study has recently described by Yasir, et al. ^33^. The transposon used to construct this library incorporates an outward-transcribing IPTG-inducible promoter. This strain was chosen due to the high quality transposon mutant library available, and because it is the parent strain for the Keio collection ^38^, an extensive library of single gene knockout mutants, which could be used to phenotypically validate the findings from the TraDIS-*Xpress* data.

### Biofilm model conditions

The library was used to inoculate parallel cultures of 5 mL LB broth (without salt) with approximately 10^7^ cells. Cultures were grown in 6-well plates containing 40 sterile 5 mm glass beads per well (Sigma, 18406). Two replicates were set up, with or without 1 mM IPTG. Plates were incubated at 30 °C with light shaking for 48 hours. After 12, 24, and 48 hours of incubation, 2 mL of planktonic sample was collected from each culture and 70 beads were taken to constitute the biofilm sample. Beads were washed twice in sterile PBS and vortexed in tubes containing PBS to resuspend cells from the biofilm. Both planktonic and biofilm samples were centrifuged at 2100 x g to form pellets for DNA extraction.

### *TraDIS-*Xpress *Sequencing*

Customised sequencing libraries were prepared to identify transposon insertions following the protocol described by Yasir, et al. ^33^. In short, DNA was extracted from pellets following the protocol described by Trampari, et al. ^87^ and was fragmented using a MuSeek DNA fragment library preparation kit (ThermoFisher). Fragments containing transposons were amplified by PCR with Tn5-i5 and i7 primers customised to recognise the transposon and the MuSeek tagged ends of the DNA ^33^. Fragments between 300 and 500 bp were size selected using AMPure beads (Beckman Coulter) and nucleotide sequences were generated using a NextSeq 500 and a NextSeq 500/550 High Output v2 kit (75 cycles) (Illumina).

### Informatics

Fastq files were aligned to the *E. coli* BW25113 reference genome (CP009273) using the BioTraDIS (version 1.4.3) software suite ^88^ using SMALT (version 0.7.6). This generated plot files for visualisation of the transposon insertion locus and frequency to compare planktonic and biofilm conditions. Insertion frequencies per gene for each replicate were plotted against each other to determine the experimental error between replicates as well as differences in insertion frequency between control and test conditions (supplementary figure 1). The tradis_comparison.R command (also part of the BioTraDIS toolkit) was also used to determine significant differences (*p* < 0.05, after correction for false discovery) in insertion frequencies per gene between control and test conditions. For all candidate loci, plot files generated by BioTraDIS were also examined manually in Artemis (version 17.0.1) ^89^ to confirm the results from these two approaches, as well as to identify regions where inserts were under differential selection but did not fall within coding regions of the genome.

### Validation experiments

Knockout mutants for genes identified by TraDIS-*Xpress* data were sourced from the Keio collection ^38^ and tested for their biofilm-forming abilities. The Keio collection is comprised of two knockout mutants per gene to account for errors in the production process, therefore each mutant was treated independently to account for any variation arising from this ^27,38^. Crystal violet assays, used to assess biofilm biomass production, were undertaken by inoculating 10^4^ of each mutant strain into 200 μL LB broth without salt in a 96-well polystyrene plate. After 48 hours incubation at 30 °C, the culture was removed, wells were rinsed with water, and the residual biofilms were stained for 10 minutes with 200 μL 0.1% crystal violet. The plate was then rinsed with water to remove the stain and 200 μL 70% ethanol was added to the wells to solubilise the stained biofilm. The optical density (OD) was measured using a FLUOstar Omega plate reader (BMG Labtech) at 590 nm. Cell aggregation was measured by leaving bacterial cultures (normalised to an OD 600 nm of 3.0) on an unagitated surface at room temperature. After 24 hours, the supernatant of each culture was removed by pipetting, diluted in PBS and measured in a plate reader at 600 nm. Biofilm matrix composition was investigated through spotting 10 μL of each mutant (representing 10^5^ CFU) on agar supplemented with 40 μg/mL Congo red (Sigma, C6277) to examine curli production. Plates were incubated at 30°C for 48 hours and photographed to compare mutant biofilm composition to the wild type. Adhesion and biofilm architecture were investigated under flow conditions for selected mutants using the Bioflux system. Flow cells were primed with LB broth without salt at 5 dyne/cm^2^ and seeded with approximately 10^7^ cells. The plate was left at room temperature for 2.5 hours to allow attachment, and subsequently incubated at 30 °C at a flow rate of 0.3 dyne/cm^2^. After 12, 24 and 48 hours, biofilms were visualised with an inverted light microscope and representative images at 10x, 20x and 40x magnification were taken at three locations of the flow cell. Experiments were performed in duplicate.

## Data availability

Sequence data supporting the analysis in this study has been deposited in ArrayExpress under the accession number E-MTAB-9873.

## Code availability statement

All software packages used are described in the methods. No bespoke code was used in this study.

## Acknowledgements

ERH was supported by a studentship funded by the Quadram Institute Bioscience. The authors gratefully acknowledge the support of the Biotechnology and Biological Sciences Research Council (BBSRC); AKT, MY, JW, IGC and MAW were supported by the BBSRC Institute Strategic Programme Microbes in the Food Chain BB/R012504/1 and its constituent project BBS/E/F/000PR10349.

## Author contributions

ERH designed and performed experiments, analysed the data and wrote the paper. AKT and IGC helped design experiments and wrote the paper. MY helped design experiments. JW heled analyse data and wrote the paper. MAW designed the experiments, analysed the data and wrote the paper.

