## Supplementary data for "Massively parallel transposon mutagenesis identifies temporally essential genes for biofilm formation in *Escherichia coli*" (from another version of this preprint: version 1, 2020-12-14)

### Supplementary Information

**Supplementary figure 1:** Insertion frequency per gene for biofilm conditions compared to planktonic conditions for each time point. Points represent the mean insertion frequency per gene for both replicates and both concentrations of IPTG.

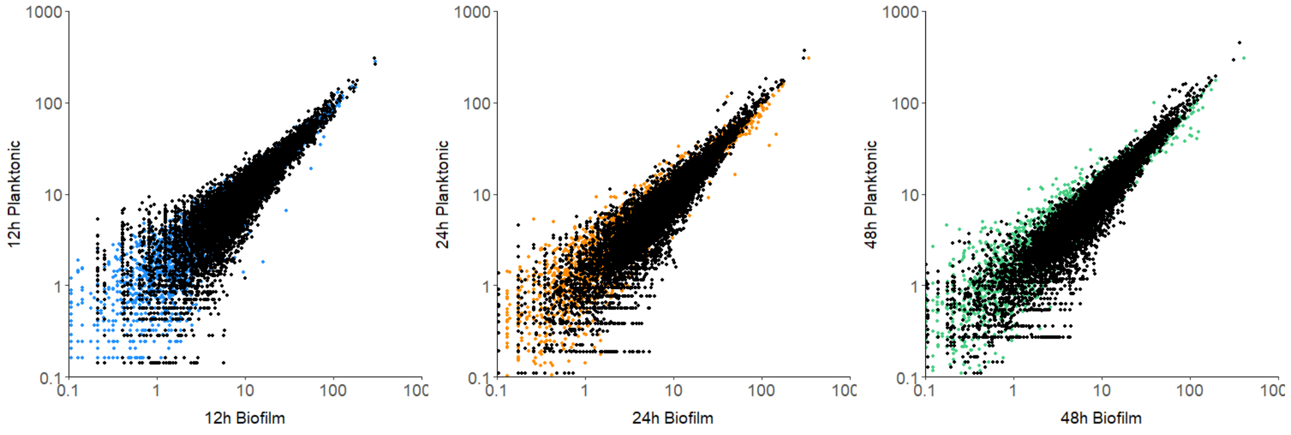

**Supplementary table 1:** Genes determined by TraDIS-Xpress to be important for biofilm formation in *E. coli*

| Pathway | Gene | Time point | Observed change in insertions in biofilm conditoion | Ref |
|---|---|---|---|---|
| Carbohydrate metabolism and transport | *sgbE* | 48h | Fewer insertions | [1] |
| Cell division | *yhcM* | 48h | Fewer insertions | [2] |
| c-di-GMP phosphodiesterase | *yfgF* | 48h | Fewer insertions | [3] |
| Curli biosynthesis | *csgC* | 24h, 48h | Fewer insertions | [4] |
| | *csgD* | 24h, 48h | Overexpression at 24h, Fewer insertions at 48h | |
| | *csgE* | 12h, 48h | Fewer insertions | |
| | *csgF* | 48h | Fewer insertions | |
| DNA housekeeping | *dam* | 24h | Fewer insertions | [5] |
| | *crfC* | 48h | Fewer insertions | [6] |
| | *maoP* | 24h | More insertions | [7] |
| Fatty acid transport | *fadL* | 48h | Fewer insertions | [8] |
| Flagella-associated motility | *flhD* | 24h, 48h | Overexpression | [9] |
| | *flhC* | 48h | Fewer insertions | |
| | *flgD* | 24h | Fewer insertions | [10] |
| | *fliE* | 48h | Fewer insertions | |
| | *lrhA* | 12h, 24h, 48h | More insertions | [11] |
| | *hdfR* | 12h, 24h | More insertions | [12] |
| LPS | *wzzB* | 48h | Fewer insertions | [13] |
| Oxidised protein repair | *yedZ* | 48h | Fewer insertions | [14] |

| | | | | |
|---|---|---|---|---|
| Outer membrane porins and channels | *mscL* | 48h | Fewer insertions | 15 |
| | *tolC* | 48h | Fewer insertions | 16 |
| | *ompF* | 48h | Fewer insertions | 17 |
| Protein modification | *dsbA* | 12h, 24h | Fewer insertions | 18 |
| Purine ribonucleotide biosynthesis | *purD* | 48h | Fewer insertions | 19 |
| | *purE* | 48h | Fewer insertions | |
| | *purH* | 48h | Fewer insertions | |
| | *purL* | 48h | Fewer insertions | |
| rRNA methyltransferase | *rlmI* | 12h | Fewer insertions | 20 |
| RNase III and biofilm regulator | *ymdB* | 24h, 48h | Fewer insertions | 21 |
| Toxin-antitoxin system | *tomB* | 12h, 24h, 48h | Fewer insertions | 22 |
| Transcriptional regulator | *dksA* | 12h, 24h | More insertions | 23-25 |
| | *leuO* | 12h, 48h | Overexpression at 12h, Fewer insertions at 48h | 26 |
| | *marR* | 12h | Fewer insertions | 27 |
| | *ompR* | 24h, 48h | Fewer insertions | 17 |
| | *lrp* | 48h | Fewer insertions | 28 |
| | *gadW* | 48h | Fewer insertions | 29 |
| | *rcsC* | 48h | Fewer insertions | 30 |
| tRNA modification | *truA* | 24h, 48h | Fewer insertions | 31 |
| Type 1 fimbriae | *fimB* | 12h, 24, 48h | Fewer insertions | 32 |
| | *fimE* | 12h, 24, 48h | More insertions | |
| | *fimC* | 48h | Fewer insertions | 33 |

| | | | | |
|---|---|---|---|---|
| | *fimD* | 24h, 48h | Fewer insertions | [33] |
| Putative fimbrial-like protein | *ydeR* | 48h | Fewer insertions | [34] |
| Unknown | *yigZ* | 12h | Fewer insertions | [35] |
| | *yebB* | 48h | Fewer insertions | [36,37] |
| | *yedN* | 48h | Fewer insertions | [38] |
| | *yjbL* | 48h | Fewer insertions | [20] |
| | *ykgJ* | 12h | Reduced expression (increased expression of antisense mRNA) | [39] |
